# Structural basis of SARS-CoV-2 spike protein induced by ACE2

**DOI:** 10.1101/2020.05.24.113175

**Authors:** Tomer Meirson, David Bomze, Gal Markel

## Abstract

**Motivation:** The recent emergence of the novel SARS-coronavirus 2 (SARS-CoV-2) and its international spread pose a global health emergency. The viral spike (S) glycoprotein binds the receptor (angiotensin-converting enzyme 2) ACE2 and promotes SARS-CoV-2 entry into host cells. The trimeric S protein binds the receptor using the distal receptor-binding domain (RBD) causing conformational changes in S protein that allow priming by host cell proteases. Unravelling the dynamic structural features used by SARS-CoV-2 for entry might provide insights into viral transmission and reveal novel therapeutic targets. Using structures determined by X-ray crystallography and cryo-EM, we performed structural analysis and atomic comparisons of the different conformational states adopted by the SARS-CoV-2-RBD.

**Results:** Here, we determined the key structural components induced by the receptor and characterized their intramolecular interactions. We show that κ-helix (also known as polyproline II) is a predominant structure in the binding interface and in facilitating the conversion to the active form of the S protein. We demonstrate a series of conversions between switch-like κ-helix and β-strand, and conformational variations in a set of short α-helices which affect the proximal hinge region. This conformational changes lead to an alternating pattern in conserved disulfide bond configurations positioned at the hinge, indicating a possible disulfide exchange, an important allosteric switch implicated in viral entry of various viruses, including HIV and murine coronavirus. The structural information presented herein enables us to inspect and understand the important dynamic features of SARS-CoV-2-RBD and propose a novel potential therapeutic strategy to block viral entry. Overall, this study provides guidance for the design and optimization of structure-based intervention strategies that target SARS-CoV-2.

## Introduction

Several members of the coronavirus family circulate in the human population and usually manifest mild respiratory symptoms (Su, et al., 2016). However, over the past two decades, emerging coronaviruses (CoV) have raised great public health concerns worldwide. The highly pathogenic severe acute respiratory syndrome-related coronavirus (SARS-CoV) (Drosten, et al., 2003; Ksiazek, et al., 2003) and Middle-East respiratory syndrome coronavirus (MERS-CoV) (Zaki, et al., 2012) have crossed the species barrier and cause deadly pneumonia in afflicted individuals, SARS and MERS, respectively. SARS-CoV first emerged in humans in Guangdong province of China in 2002, and its global spread was associated with 774 deaths among 8,096 cases (de Wit, et al., 2016; WHO, 2004). In 2012, MERS-CoV was first identified in the Arabian Peninsula and spread to 27 countries, infecting a total of 2,494 people and claiming 858 lives (WHO, 2020). While the SARS pandemic was finally stopped by conventional control measures, including patient isolation and travel restrictions, new cases of MERS have been reported (Yoon and Kim, 2019).

In December 2019, a previously unknown CoV, named SARS-CoV-2, was discovered in Wuhan, Hubei province of China (Huang, et al., 2020; Wang, et al., 2020; Zhu, et al., 2020). The sudden emergence of the novel SARS-CoV-2 has rapidly evolved into a pandemic that posed a serious threat to global health and economy (Gates, 2020). Although SARS and MERS have a higher mortality rate, SARS-CoV-2 infection (COVID-19) spreads much more rapidly (To, et al., 2020). As of May 2020, more than 4,000,000 confirmed infections were reported in 216 countries, including over 300,000 deaths (WHO, 2020). MERS-CoV, SARS-CoV, and SARS-CoV2 were suggested to originate from bats and most likely serve as a reservoir host for these viruses (Ge, et al., 2013; Haagmans, et al., 2014; Li, et al., 2005; Memish, et al., 2013; Zhou, et al., 2020). Detailed investigations of the zoonotic origin of human CoVs indicate that SARS-CoV was transmitted from palm civets to humans and MERS-CoV from dromedary camels to humans (Guan, et al., 2003; Haagmans, et al., 2014; Kan, et al., 2005). However, the intermediate host for zoonotic transmission of SARS-CoV-2, linked to a wet animal market in Wuhan, is still under investigation (Walls, et al., 2020; Ye, et al., 2020).

The spike (S) glycoprotein is a class I fusion protein that mediates the entry of CoVs into target cells (Tortorici, et al., 2019). The S protein forms homotrimers that protrude from the viral surface (Figure 1A) and comprises two functional subunits, which facilitate viral attachment to the surface of host cells (S1 subunit) and fusion of the viral and cellular membranes (S2 subunit) (Figure 1B). The distal part of S1 subunit, the receptor-binding domain (RBD), is linked through two anti-parallel hinge linkers which connect the domain to the N-terminal domain (NTD) and C-terminal domain 2 (CTD2) and allow the transition between closed and open conformations (Gui, et al., 2017; Song, et al., 2018; Walls, et al., 2020; Yuan, et al., 2017) (Figure 1A-C). The open conformation of the S1 subunit facilitates interaction with angiotensin-converting enzyme 2 (ACE2) (Figure 1D-E), which contributes to the stabilization of the prefusion state of the S2 subunit that contains the fusion machinery (Song, et al., 2018; Walls, et al., 2020). Activated S protein is cleaved by host proteases at the S1/S2 and S2’ site resulting in a cleaved S2’ subunit that drives the fusion of viral and cellular membranes (Belouzard, et al., 2009; Millet and Whittaker, 2014; Pallesen, et al., 2017). The critical step in SARS-CoV-2 infection which involves the transition between a metastable prefusion state to a stable post-fusion state is triggered by binding to ACE2 which induces conformational changes in the RBD and the hinge region (Gui, et al., 2017; Pallesen, et al., 2017; Song, et al., 2018; Walls, et al., 2020). While the interaction between the S protein and ACE2 has been extensively studied (Han, et al., 2006; Lan, et al., 2020; Letko, et al., 2020; Li, 2013; Shang, et al., 2020; Song, et al., 2018; Wan, et al., 2020; Wang, et al., 2020; Yan, et al., 2020), the key determinants in the activation of the virus upon binding to the host receptor is poorly understood. We aimed to investigate the structural basis of the SARS-CoV-2 S protein modulation induced by virus-receptor interaction.

**Figure 1.**
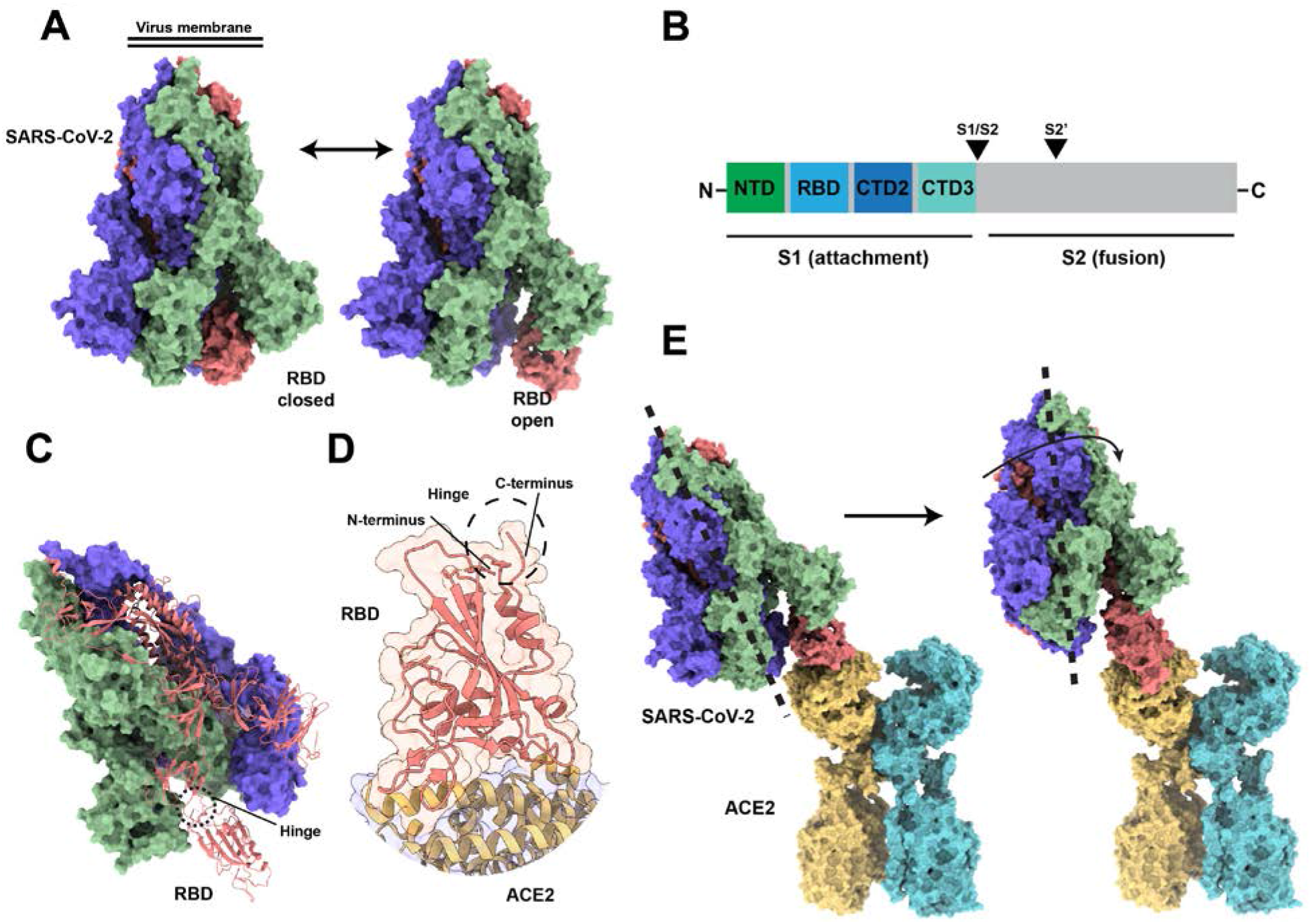
Structural comparison of SARS-CoV-2 S protein conformational states. (A) Surface diagram of SARS-CoV-2 homotrimeric structure in the unbound-closed and open conformations. (B) Structural illustration of S protein, including functional domains (NTD, N-terminal domain; RBD, receptor-binding domain; CTD2, C-terminal domain 2; CTD, C-terminal domain 3; and proteolytic cleavage sites (S1/S2, S2’). (C) S trimer with one RBD in the open conformation and (D) RBD-ACE2 complex shown as a cartoon. (E) Superposed structures depicting the conformational changes between the unbound-open (left) to the ACE2-bound state.

## Methods

### Structure collection and modeling

We searched the PDB database for high-quality structures of the SARS-CoV-2 S protein at closed, open, and bound conformations. For each conformation, we used the structure with the lowest resolution that contains the RBD and the hinge region. For the unbound-closed, unbound-open, and bound conformations, we used PDB IDs 6VXX:B, 6VYB:B, and 6M0J:E, respectively. To model the position of SARS-CoV-2 S protein compared to ACE2, we structurally aligned the models to the RBD of ACE-B^o^AT1 complex (Yan, et al., 2020) (6M17) and SARS-CoV in the bound conformation with the highest degree of RBD opening (Song, et al., 2018) (6ACK). The structures were preprocessed with Dock Prep and aligned using UCSF Chimera v1.13 (Pettersen, et al., 2004). To compare the secondary structure compositions of attachment proteins, we used structures with the lowest resolution from different families of enveloped viruses with class I fusion proteins (White, et al., 2008). These structures were compared to a random dataset comprising the first 1000 non-viral PDB proteins deposited in 2020 with a resolution <3 Å.

### Secondary structure assignment

The initial assignment of secondary structure was performed using DSSP (Touw, et al., 2015). To assign κ-helix (alternative designation for polyproline II [PPII]), we used the method which was recently introduced (Meirson, et al., 2020; Meirson, et al., 2020). Briefly, we calculated the root-mean-square dihedral deviations (RMSdD) of the peptide backbone torsional angles φ and Ψ as a measure of the average deviation from a reference κ-helix. To include short segments of κ-helix, at least two consecutive residues with mean RMSdD below the cutoff (ε) of 17 (Mansiaux, et al., 2011) were defined as the criteria for the assignment.

### Structural characterization

To analyze the conformational changes induced by ACE2 binding, we compared the bound with the unbound form. We included both the open and closed states of the unbound structures to highlight the specific effects induced by the interaction with ACE2. The backbone dihedral angles (φ, Ψ) were converted to generic helix parameters ϑ (angular step per residue) and d (rise per reside) as described by Miyazawa (Miyazawa, 1961) to describe the geometrical variations intuitively. Van der Waals (VDW) intramolecular interactions were determined using a distance cutoff of 3.5 Å. Only interactions that were gained or lost by the bound compared to both unbound conformations were considered. To identify interacting residues with considerable shifts, displacements smaller than 0.5 Å were excluded. To identify important backbone hydrogen bonds (H-bonds), the mean difference in the electrostatic interaction energy between the bound and the unbound conformations was calculated using DSSP. For each acceptor-donor pair, gain, or loss of H-bond was defined if the mean energy difference is <−1kcal/mol or >1kcal/mol, respectively. The magnitude of the energy threshold represents twice the standard cutoff (−0.5kcal/mol) for the existence of a H-bond (Zhang and Sagui, 2015). Structural analyses was performed with the Bio3D package (Grant, et al., 2006) in R version 3.6.

### Disulfide bond analysis

Due to inconsistencies in the reported composition of disulfide bonds in the RBD in different structures (Lavillette, et al., 2006; Song, et al., 2018; Walls, et al., 2020; Wrapp, et al., 2020; Yuan, et al., 2017) and to account for possible errors in modeling disulfide bonds (Carpentier, et al., 2010; Kleywegt and Jones, 1995; Villa and Lasker, 2014; Wlodawer, et al., 2008), we used Disulfide by Design 2.0 (DbD2) to calculate χ3 torsion angles and bond energies (Craig and Dombkowski, 2013). The DbD2 algorithm could accurately predict the chiralities and positions of disulfide bonds based on energy function, reflecting the geometric characteristics of disulfide bonds among high-quality crystal structures (Craig and Dombkowski, 2013; Wiedemann, et al., 2020). We used the estimated disulfide energy threshold of <2.17kcal/mol that applies to most naturally occurring disulfides, χ3 angles of −87 or 97 ± 20 (Craig and Dombkowski, 2013) and disulfide bond distance of 2.03 ± 5% (Spek, 1990) to indicate a high probability of a disulfide bond that satisfies stereochemical constraints.

### Graphical visualization

The models were visualized using UCSF ChimeraX v0.94 (Goddard, et al., 2018). Models with reassigned secondary structures were visualized using the academic version of Schrodinger Maestro v11.1 (Bell, et al., 2006). κ-helices and 3_10_-helices were represented as ribbons and tubes, respectively. Ribbons were drawn, passing through carbon alphas. Trajectories were produced by interpolating between the bound and unbound-closed conformation and visualized using VMD (Humphrey, et al., 1996).

## Results

### Secondary structure determination

To characterize the structural composition of the RBD of SARS-CoV-2 and other members of the class I fusion proteins, we calculated the distribution of secondary structure assignment. Since κ-helix (PPII) often serves functional purposes in proteins (Adzhubei, et al., 2013; Meirson, et al., 2020) and DSSP does not assign the conformation, we performed reassignment of the secondary structures to reveal potential κ-helix conformations. Compared to a random dataset in the PDB, structures of the attachment proteins of CoVs, influenza, measles, HIV and Ebola viruses display higher proportions of β-strands and κ-helices, whereas α-helix is under-represented (Figure 2A). Among non-regular secondary structures, the random coil is the most over-represented assignment. The reassigned SARS-CoV-2 structure reveals a diverse distribution of κ-helices throughout the domain, whereas other secondary structures are clustered more closely together (Figure 2B-C). The most common secondary structure in the ACE2 binding interface is κ-helix (Figure 2C).

**Figure 2.**
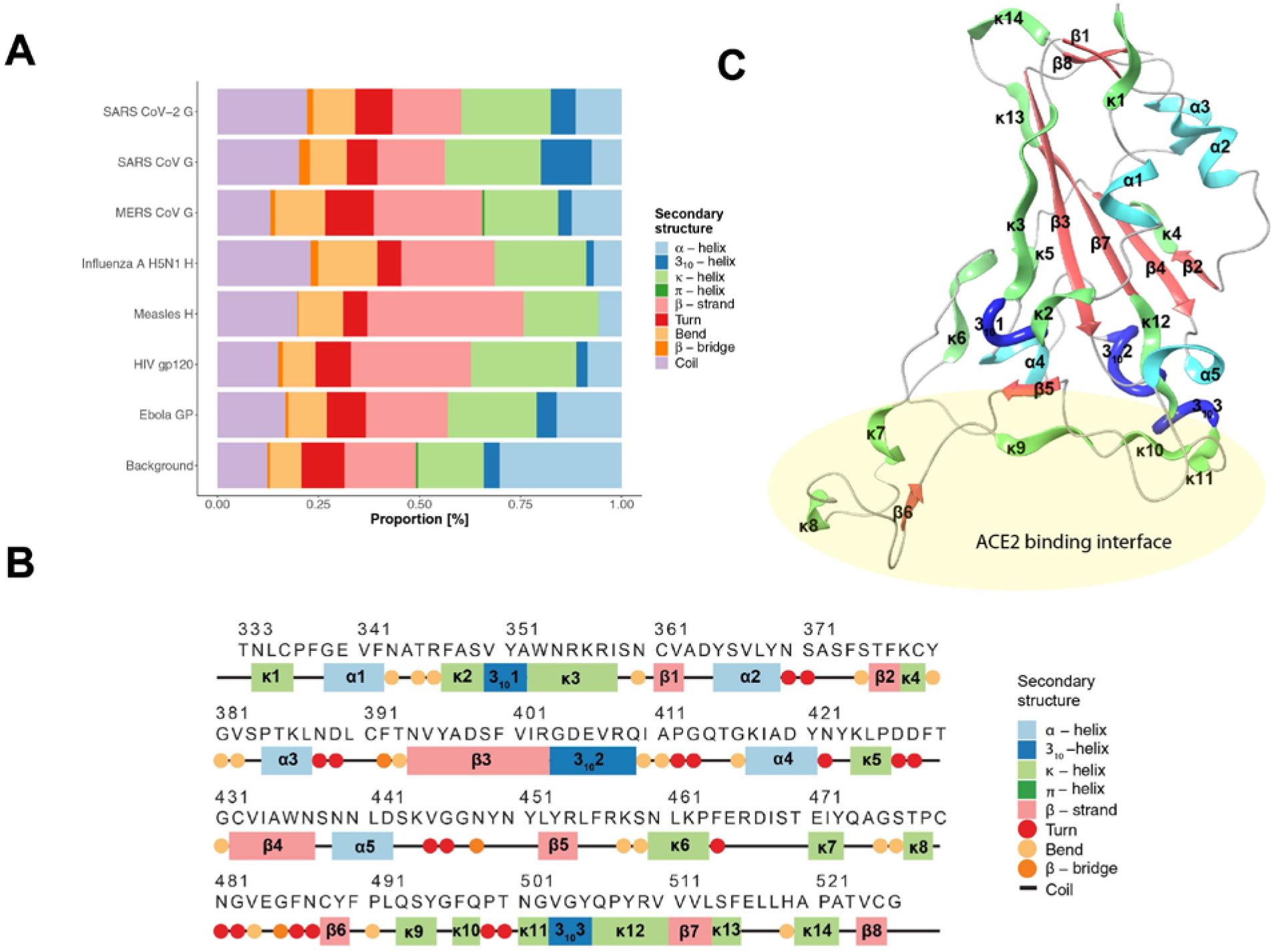
Secondary structure assignment of the receptor-binding domain. (A) Comparison of attachment proteins of coronaviruses and other members of the class I fusion proteins. Position of secondary structure assignment (B) and cartoon representation (C) of SARS-CoV-2-RBD. The secondary structural elements are labeled according to their occurrence in sequence. α-helices (cyan), β-strands (red), and κ-helices (green) are illustrated as ribbons and 3_10_-helices (blue) as thick tubes.

### Structural variations of SARS-CoV-2-RBD bound with ACE2 receptor

To gain insights into the effects of ACE2 interaction on the S protein of SARS-COV-2, we analyzed the intramolecular structural variations in the bound versus the unbound-closed and unbound-open conformations. Figure 3 depicts a summary of the main differences in the intramolecular interaction and H-bond profile upon receptor binding and allows to follow the interconnectivity path leading to the hinge region at the termini. While small changes in the rotational angles or rise of residues occur throughout the domain (Figure 3A), the most substantial changes happen at non-regular secondary structures such as coils and turns. Compared with the unbound-open state, the bound conformation is mainly associated with the formation of κ-helices (20 residues), followed by α-helices (13 residues), β-strands (10 residues), and a 3_10_-helix (3 residues). The interaction map demonstrates a rich rearrangement of intramolecular interactions (Figure 3A). Interactions are gained mostly in missing loops at the binding interface (455-491), whereas lost interactions are scattered throughout the domain. Figure 3B shows the redistribution of main-chain H-bonds energy and rearrangement of acceptor and donor H-bond networks. Consistent with the interconnectivity map, the pairing of H-bonds occurs at the ACE2 binding interface. Interestingly, local rearrangements of H-bonds occur mainly at α-helices, whereas κ-helices facilitate distant H-bonds. This indicates that κ-helices mediate switch-like interactions.

**Figure 3.**
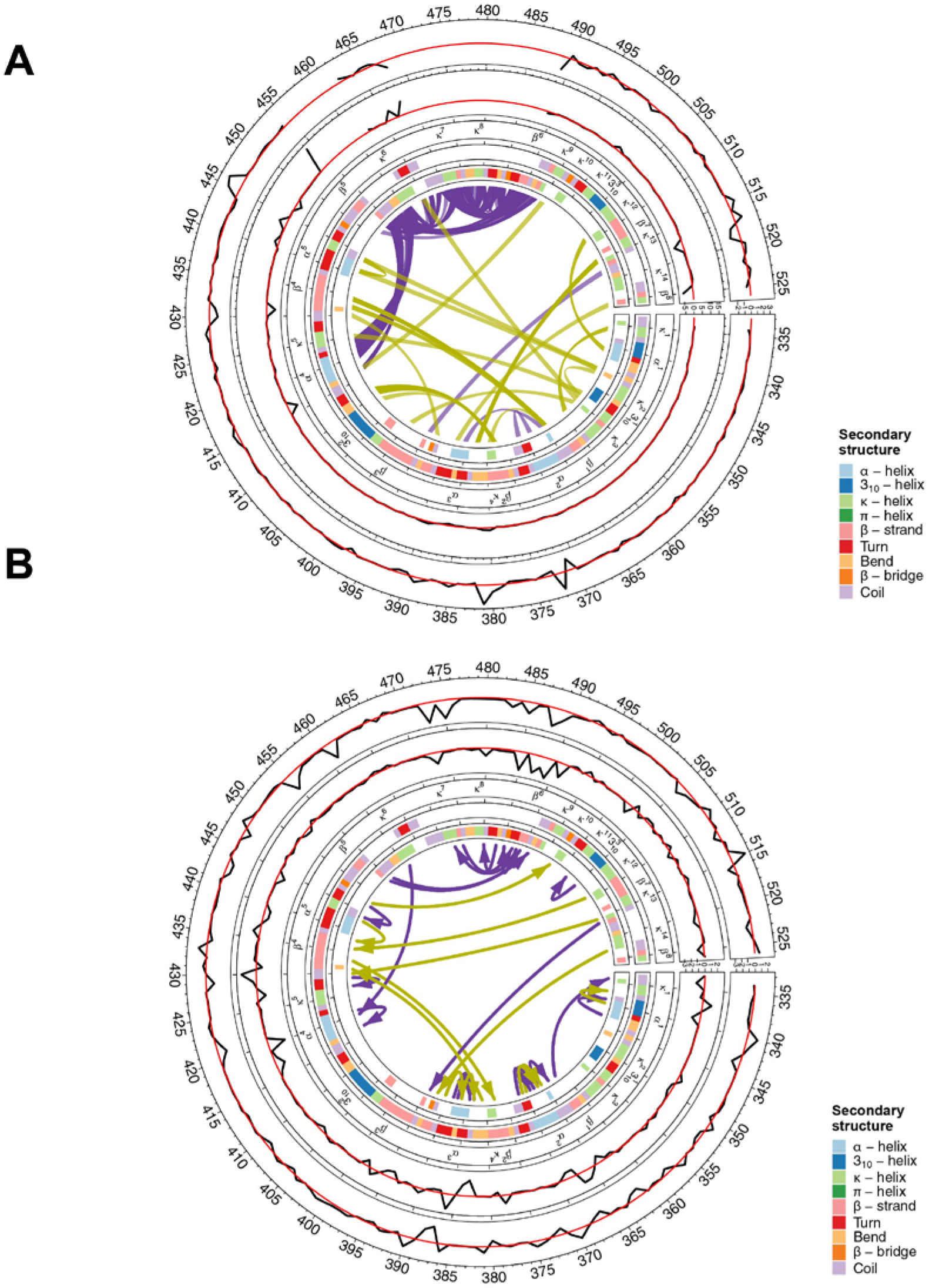
Intramolecular interactions induced by SARS-CoV-2-RBD and ACE2 complex. (A) Circos plot depicting the main differences in intramolecular interactions (< 3.5Å) between the unbound (6VXX:B and 6VYB:B) and bound (6M0J:E) conformations of the SARS-CoV-2-RBD. Track order: shift in the angular step (ϑ) per residue (i); shift in the rise (d) per residue (ii); labels of secondary structural elements (iii); secondary structure assignment of the unbound-open conformation (iv); transformed secondary structure of the bound conformation, representing the induced conformational changes (v). Gained and lost interactions are shown as magenta and yellow lines, respectively. (B) Circos plot depicting the main differences in main-chain H-bond profile between the unbound and bound conformations of the SARS-CoV-2-RBD. Track order: mean difference in the electrostatic interaction energy of the donor (i) and acceptor (ii); labels of secondary structural elements (iii); secondary structure assignment of the unbound-open conformation (iv); transformed secondary structure of the bound conformation, representing the induced conformational changes (v). Gained and lost H-bonds (|ΔE| > 1kcal/mol) illustrating the direction of donor-acceptor are shown as magenta and yellow arrows, respectively.

A closer inspection into the structural variations induced by the interaction SARS-CoV-2-RBD and ACE2 is shown in Figure 4. Upon binding, three H-bond networks comprising residues V504, Y505 with Q506, D442, Y495 with F497, and D442, N448 with K444, are destabilized (Figure 4A). The loss of H-bonds of Y495 and the side chains of K443 and Y505 is associated with conversion between κ5’ into a coil and the formation of a new κ-helix κ10. This results in the advancement of D442 and the stabilization of α-helix α5. The amino-acid R509 is rotated and switches between a β-strand (β7) to a κ-helix (κ12) while gaining a solid network of H-bonds and salt bridge with the side chain of D442 in the newly formed α5, at the expense of losing H-bonds with the backbone of V341 and F342 (Figure 4B). Consequently, a new H-bond is formed between N343 and G339, which converts 3_10_1’ into α1. The adoption of a shorter-pitched α-helix pulls κ1, which constructs the N-terminus of the hinge region. The main-chain H-bond of F515 with G314 and the π-π interaction with F392 are lost (Figure 4C) while a stronger pairing of H-bond forms between G431 and Y380 (Figure 3B). The rearrangement of the interactions is associated with conformational changes, including the conversion between κ14’ to β8, which constructs the C-terminus of the hinge region. Also, the loop downstream to Y380 is displaced, and α-helix α3 is formed by a contribution of H-bond between the backbones of P384 and N388. The H-bond between L397 and 390 is lost, and the new α3 packs closely together with α2, which includes VDW interactions between S366 and T385 (Figure 4D). Notably, this interaction is among the only gained contacts observed in the RBD, not involving the missing loops (Figure 3A). Also, a concentration of H-bonds is re-distributed along α2, repositioning the α-helix one step back (from 366-370 to 365-369). This transition is associated with a small movement of β1 at the medial region of the hinge.

**Figure 4.**
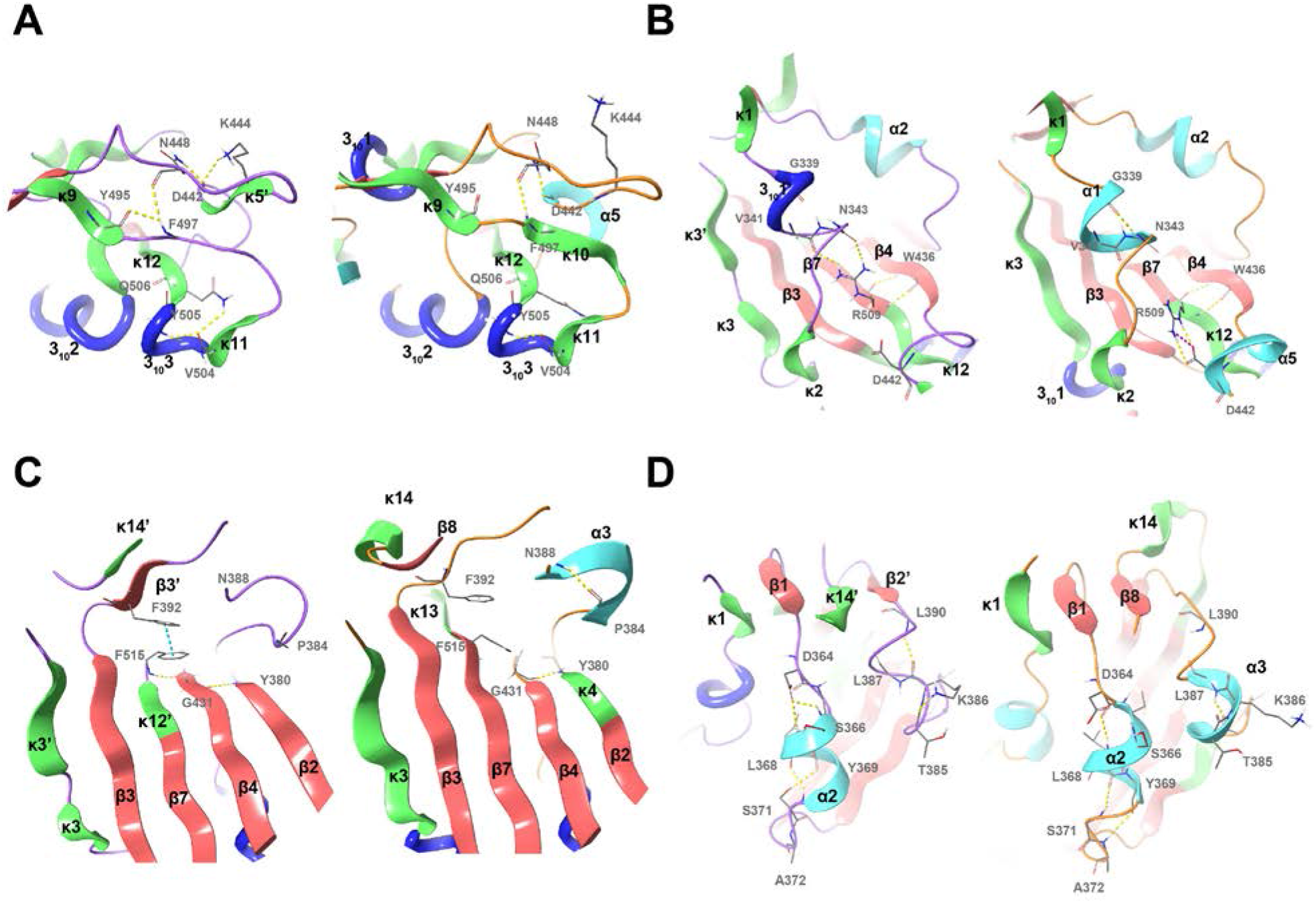
Detailed structural comparison between the unbound and bound SARS-CoV-2-RBD and ACE2. (A-D) Representative structures comparing the unbound-open (6VYB:B) and bound (6M0J:E) conformations are shown as cartoon representation. Key contacts are labeled and shown as sticks. The secondary structural elements are labeled according to their occurrence in sequence. α-helices (cyan), β-strands (red), and κ-helices (green) are illustrated as ribbons and 3_10_-helices (blue) as thick tubes.

### Dissociation between the RBD and CTD2

The κ13/κ14 loop between residues 515 and 523 exhibits a relatively large conformational change between the unbound-closed and the bound states (Supplementary Figure 1). This loop is involved in the interaction between the RBD and CTD2 in the closed conformation. However, in the open conformation where RBD dissociates from CTD2, the loop is missing in both SARS-CoV and SARS-CoV-2 (Song, et al., 2018; Walls, et al., 2020; Wrapp, et al., 2020). The dissociation between the domains is required for the hinge and the RBD to move freely. Together, this indicates that the RBD-ACE2 complex stabilizes the κ13/κ14 loop in a conformation that disfavors the interaction between RBD and CTD2.

### Disulfide bond analysis

The RBD-ACE2 complex inducible interactions culminate in the hinge region. The hinge, which we show to exhibit various conformational changes (Figure 3 and 4), contains two pairs of cysteines. Since cysteine pairs have the potential to act as allosteric switches (Bekendam, et al., 2016; Butera, et al., 2018; Chiu and Hogg, 2019), we hypothesized they are altered during the interaction between the S protein and ACE2 receptor. Therefore, we calculated the energy and geometrical features of the potential disulfide bonds in all high-quality structures of the RBD in SARS-CoV-2. Due to gross inconsistencies between disulfide assignments (Lavillette, et al., 2006; Song, et al., 2018; Walls, et al., 2020; Wrapp, et al., 2020; Yuan, et al., 2017) and quality issues of these pairs, we also characterized quality metrics of the cysteine pairs. Figure 5A shows an alternating pattern in the quality metrics of the N-(Cys^336^-Cys^361^) and C-terminal (Cys^391^-Cys^525^) cysteine pairs, except one structure (PDB 6VYB:B) in which the pair Cys^336^-Cys^361^ in the open conformation is in the reduced form. This indicates that only one disulfide exists at a given state. Among cysteine pairs without assignment issues, the disulfide bond energy also shows an alternating pattern (Figure 5B), where the unbound-open and bound conformation have low energies in the N- and C-terminal cysteine pairs, respectively. This indicates that the cysteine pair switch between disulfide classes upon binding of the RBD to the receptor. Since the two cysteine pairs are aligned parallelly and closely together, the results suggest they are involved in disulfide shuffling (Figure 5B,C).

**Figure 5.**
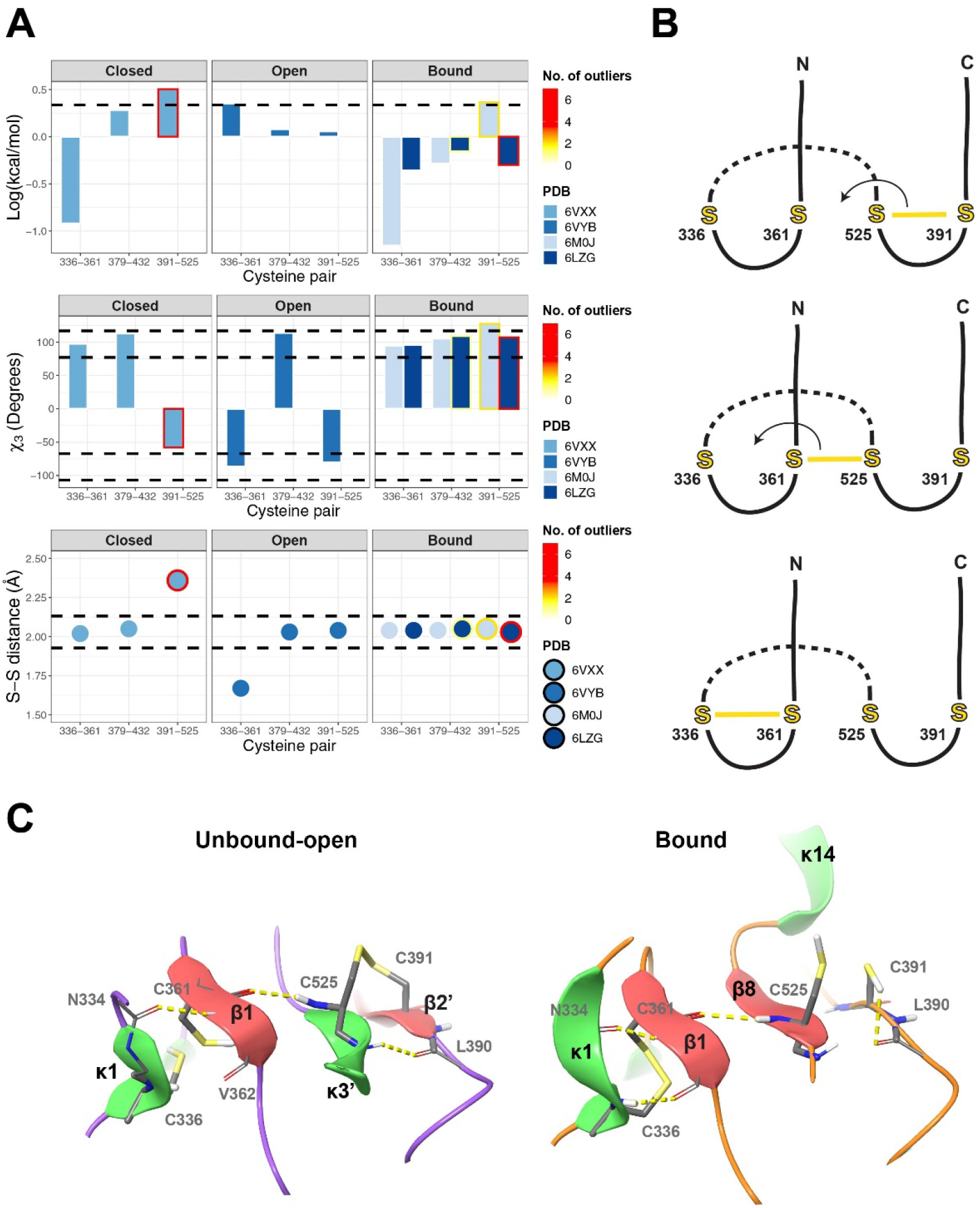
Characterization of the conserved SARS-CoV-2-RBD cysteine pairs. (A) Shown are the disulfide bond energy (top), χ3 torsional angle (middle), and S-S distance (bottom) in different conformational states of SARS-CoV-2-RBD (closed – 6VXX:B; open – 6VYB:B; bound – 6M0J:E and 6LZG:B). Dashed lines show the threshold for ideal disulfide bonds (top panel – below the line; middle and bottom panel – between the lines). The number of outliers, including RSRZ (root-mean-square of Z score) outliers, non-rotameric sidechains, clashes, and bond length and angle outliers, were summarized for each pair. Of note, the disulfide outliers were among the highest-ranked outliers of all the corresponding structure, indicating a poor fit to the density map. (B) Illustration of the predicted disulfide shuffling between the conserved four cysteine residues. (C) A cartoon representation of the hinge region with the two cysteine pairs depicting alternating disulfide bond configurations. The unbound-open conformation (left) demonstrates the original disulfide bond configuration. In the bound conformation (right), the Cys^391-525^, which poorly fit the density map, are shown as unpaired cysteines.

A summary of the conformational changes between the unbound and the bound RBD states are shown as an interpolated trajectory in Figure 6A. Also, a simplified model linking between the distal part and the hinge is depicted in Figure 6B.

**Figure 6.**
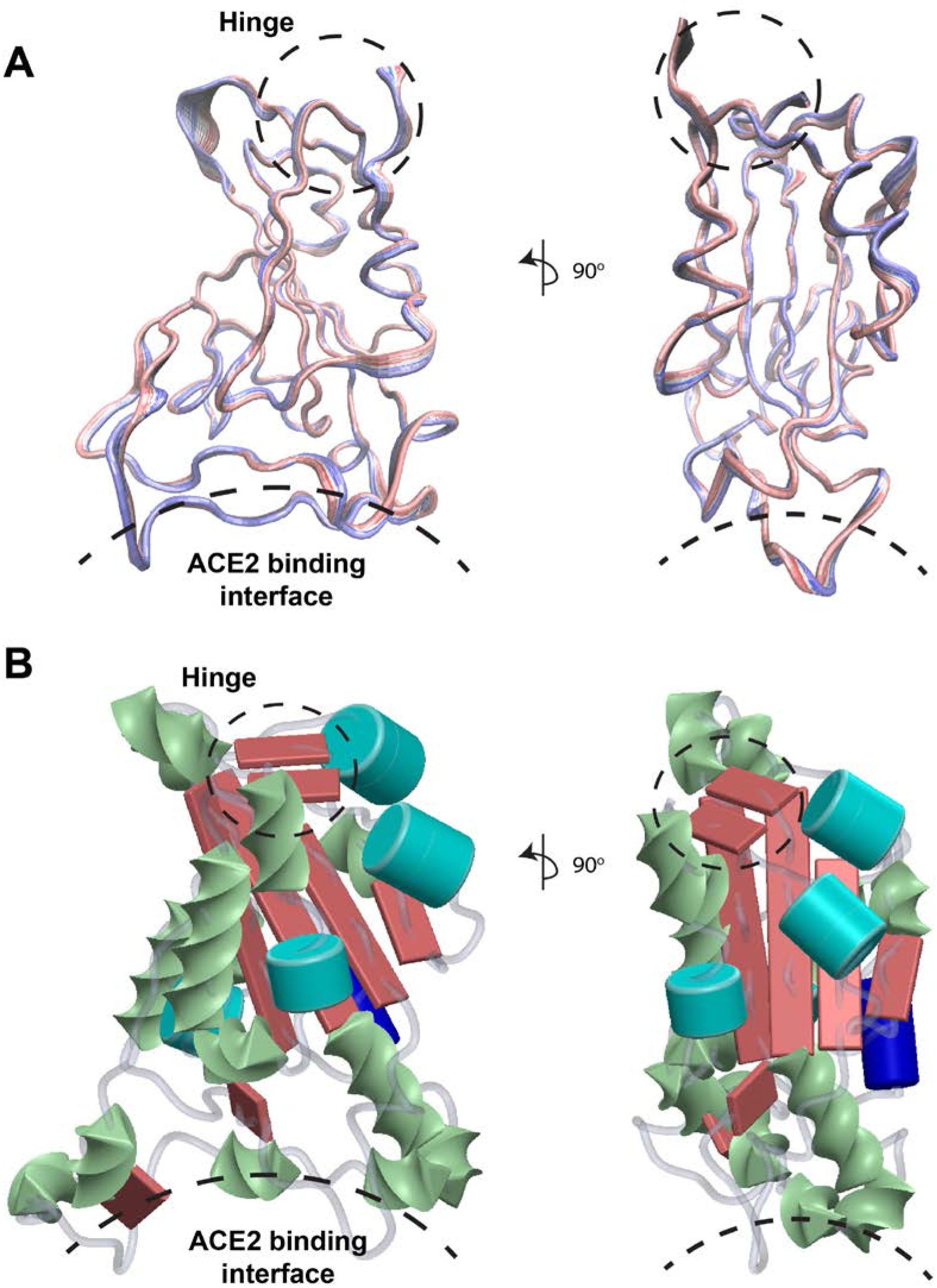
Dynamical features of the SARS-CoV-2-RBD. A summary of the conformational changes between the unbound-closed (6VXX:B in blue) and the bound (6M0J:E in red) RBD states are shown as an interpolated trajectory. (B) A simplified model, demonstrating the association between the secondary structures and the relationship between the distal part (ACE2 binding interface) and the hinge region. α-helices (cyan), β-strands (red), and κ-helices (green) are illustrated as shown as 3D shapes.

## Discussion

The recent emergence of SARS-CoV-2 pandemic represents a major epidemiological challenge. ACE2 has been reported to be the receptor that initiates the activation of this novel CoV (Hoffmann, et al., 2020; Yan, et al., 2020). In this study, we determined the key structural components induced by the receptor and characterized their intramolecular interactions.

Numerous structures of the prefusion human CoV S proteins were determined at different states, and the key regions responsible for the interaction with the receptor were previously reported (Lan, et al., 2020; Shang, et al., 2020; Walls, et al., 2020; Wan, et al., 2020; Wrapp, et al., 2020; Yan, et al., 2020). However, to our knowledge, no previous study has extensively investigated the mechanism of transduction through the RBD. The structural transduction mechanism on a molecular level remains a difficult question to address experimentally. Therefore, we used current state-of-the-art structures of the SARS-CoV-2-RBD and focused on their structural organization. These structures represent snapshots of the dynamic S protein and allow to track and model the conformational transitions between the different states.

Characterizing the secondary structures of proteins is fundamental for gaining knowledge and simplifying the complicated 3D structures. We show that κ-helix is a predominant structure in the binding interface and in facilitating the conversion to the active form of the S protein. This conformation which is commonly known as PPII was recently designated as κ-helix, following the widespread criticism of the misleading name PPII (Adzhubei, et al., 2013; Hollingsworth, et al., 2009; Mansiaux, et al., 2011; Martin, et al., 2014; Meirson, et al., 2020; Meirson, et al., 2020). As many structures contain few prolines or none, the name ‘polyproline’ is considered inappropriate, and a more general term which abides the tradition of Latin letters to secondary structures was proposed (Meirson, et al., 2020). The role of κ-helix in propagating interactions and facilitating switch-like components coincides with the assessment that they represent ‘functional blocks’ as compared with other conformations such as α-helices that often represent structural building blocks (Adzhubei, et al., 2013; Meirson, et al., 2020). The flexible and extended conformation of κ-helices, as well as non-regular H-bonds and preferred location on the surface of proteins, making them ideal elements for a wide range of molecular interactions (Cubellis, et al., 2005; Stapley and Creamer, 1999; Zagrovic, et al., 2005). However, despite being more common than most secondary structures, this conformation is often overlooked, apart from proline-rich regions. This is explained in part because it is not defined by H-bonds and is not assigned by the secondary structure assignment program employed in the PDB. Other reasons include a lack of graphical representation and its misleading historical name (Meirson, et al., 2020).

Our findings demonstrate that the high prevalence of κ-helix, as well as β-strands, are not unique to SARS-CoV-2 and appear to characterize other viruses. The conformational changes between different states of the SARS-CoV-2-RBD are associated with a typical transition between β-strand and κ-helix, as they are closely related in the torsional space (Hollingsworth and Karplus, 2010; Mansiaux, et al., 2011; Oh, et al., 2010). Therefore, we hypothesize that κ-helix could serve as an efficient evolutionary tool due to its flexible nature, which could adapt more quickly in the dynamic environment compared to more restricted secondary structures. In line with this suggestion, Austin et al. showed evolutionary conservation of κ-helix bias in intrinsically disordered regions that could be tuned by changing the distribution of κ-helix for multiple functions, including molecular recognition or allosteric regulation (Austin Elam, et al., 2013).

The hinge region was reported to facilitate the RBD motion and participate in the activation process (Gui, et al., 2017; Pallesen, et al., 2017; Song, et al., 2018; Walls, et al., 2020), and our structural analysis suggests that the conformational changes culminate at the hinge which contains four highly conserved cysteines (Shang, et al., 2020; Wang, et al., 2020). To explore a possible allosteric switching mechanism, we performed atomic comparisons of the cysteine pairs at different states of the S protein. Since inconsistencies in disulfide assignment exist and errors in structure determination are not uncommon (Carpentier, et al., 2010; Kleywegt and Jones, 1995; Villa and Lasker, 2014; Wlodawer, et al., 2008), we also assessed their geometric quality. Atomic details of the structures at a pH range of 6.5-8.0 reveal alternating patterns in bond energy, geometric characteristics, and quality, between the pairs Cys^336^-Cys^361^ and Cys^391^-Cys^525^, but not in Cys^379-432^. Nonetheless, Cys^379-432^ displays a switch in the chirality of the disulfide bond in the bound state. These findings demonstrate a switching mechanism between disulfide bonds of Cys^336^-Cys^361^ and Cys^391^-Cys^525^, where at each state (open, closed, or bound), only one pair of cysteine satisfies favorable disulfide configuration and quality criteria. The unfavorable configuration is associated with unphysical disulfide bond characteristics, high energy, poor quality, or their combination. This indicates that the distorted disulfide bonds entail substantial stress or that bond assignments were inaccurate, and these pairs are reduced. Both are possible as the cysteine pairs, located at the hinge, undergo significant conformational changes, and forced stretching of disulfide bonds is known to accelerate their cleavage (Zhou, et al., 2014). The four cysteine residues are adjacent and aligned suitably for disulfide exchange reactions. Such an arrangement makes it possible for a concerted series of disulfide exchange reactions to occur (Zhou, et al., 2008) and is supported by the alternating pattern of the Cys^336^-Cys^361^ and Cys^391^-Cys^525^ disulfide bond configurations. A proposed model of disulfide shuffling and SARS-CoV-2 viral entry is depicted in Figure 7.

**Figure 7.**
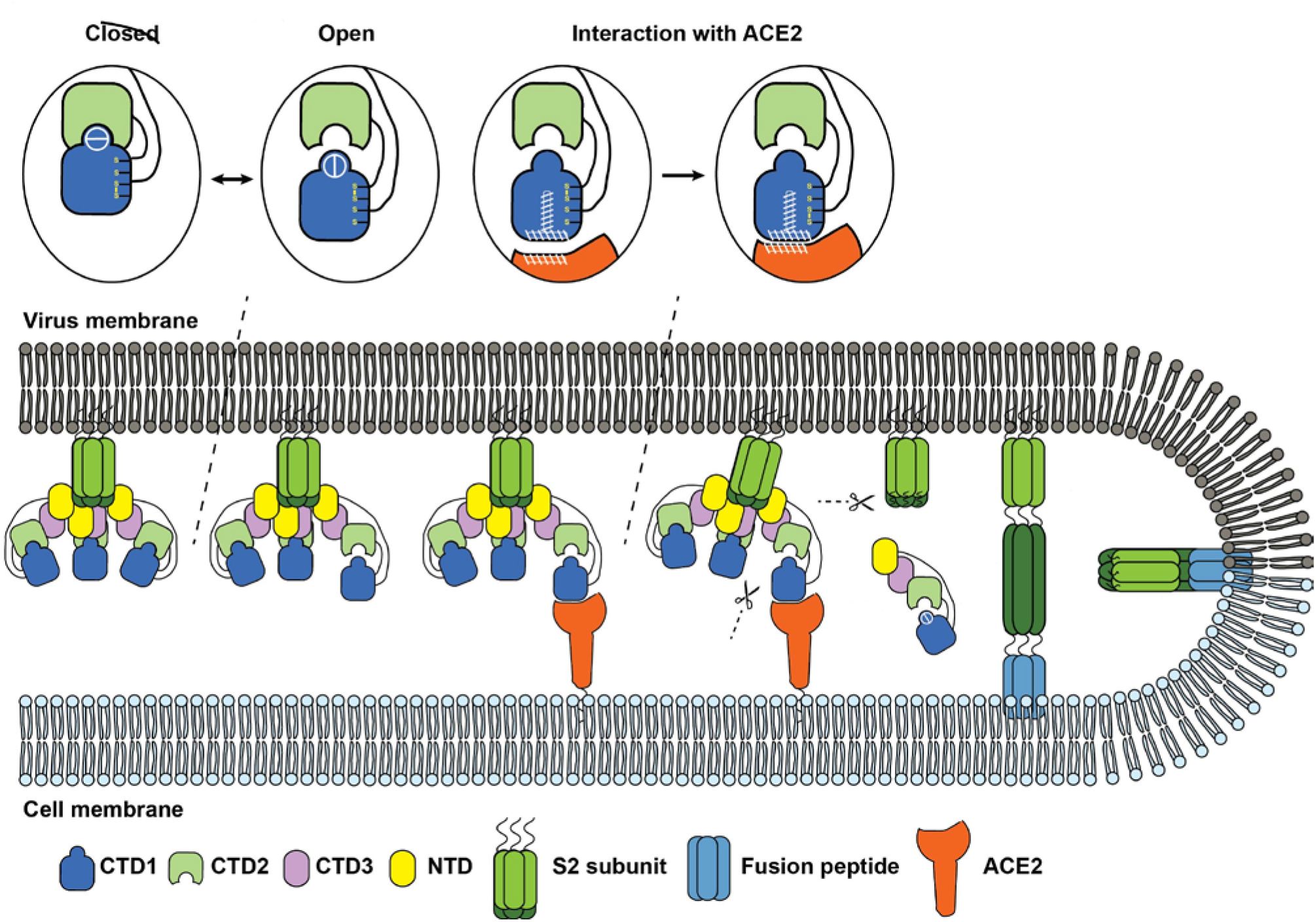
Proposed model for the pre- to post-fusion transition of SARS-CoV-2 S protein. The closed and open transitions are associated with the dissociation of RBD and CTD2 and conformational changes in the hinge region, including disulfide bond rearrangement. The complex of SARS-CoV-2-RBD with ACE2 induces conformational changes and rearrangement of the disulfide bonds that stabilize the S protein in the active form. The Activated S protein is cleaved by host proteases and induces the pre- to post-fusion transition of the S2 subunit, and initiates the fusion of viral and cellular membranes.

Beyond serving purely structural role, disulfide bonds can participate in redox reactions and act as allosteric switches controlling protein functions (Bekendam, et al., 2016; Butera, et al., 2018; Chiu and Hogg, 2019; Zhou, et al., 2014). Specific disulfide exchange reactions depend on a reducing agent such as thioredoxin or protein disulfide isomerase (PDI) (Zhou, et al., 2014). Rearrangement of disulfides (disulfide shuffling) can also occur via intra-protein thiol-disulfide exchange reactions without additional agents, which depends on conformational changes (Chiu and Hogg, 2019; Zhang, et al., 2018). An increasing number of studies support an essential role for disulfide exchange in the entry of multiple viruses in susceptible cells (Stantchev, et al., 2012). In HIV, the attachment of gp120 subunit of the viral envelope (Env) to its primary receptor CD4, induces conformational changes that cause disulfide exchange in a PDI-dependent manner and is obligatory for triggering membrane-fusion process (Fenouillet, et al., 2007; Owen, et al., 2016; Stantchev, et al., 2012). More specifically, structural rearrangements of the S protein of CoV murine hepatitis virus (MHV) during cell interaction has been reported to affect cell entry using disulfide shuffling in the RBD (Gallagher, 1996; Weismiller, et al., 1990) as our result suggest for SARS-CoV-2. Surprisingly, Lavillette et al. showed that SARS-CoV S1 subunit is redox insensitive using chemical manipulation of the redox state, in contrast to various viruses including HIV and the CoV MHV (Fenouillet, et al., 2007; Lavillette, et al., 2006). However, the study utilized murine leukemia retrovirus (MLV) pseudotyped with S1 subunit, lacking the S2 subunit, a system with limited biological relevance as the subunits cooperate and form a tightly packed trimeric structure. Furthermore, the subunits remain non-covalently bound after proteolytic S1/S2 cleavage (Tortorici, et al., 2019; Walls, et al., 2016).

We propose that targeted redox exchange between conserved cysteine pairs in the S protein could conceptualize a new strategy in the development of high-affinity ligands against SARS-CoV-2, with important therapeutic implications. Recently, Hati et al. showed, using molecular dynamic simulations, that reducing all disulfide bonds in both ACE2 and SARS-CoV2 impairs their binding affinity (Bhattacharyay and Hati, 2020). However, more evidence is required to establish the role of redox potential and paired and unpaired cysteines in the S protein during viral entry. Also, our study is limited to a computational assessment of structures reconstructed using X-ray and cryoEM, and the implications of the observed structural rearrangements remain to be determined.

Currently, no efficient antivirals against SARS-CoV-2 or other CoVs are available, and numerous clinical trials are underway (Lythgoe and Middleton, 2020). In parallel, efforts continue to develop antivirals and vaccines (Chen, et al., 2020; Liu, et al., 2020). Structure-based design of antivirals that efficiently recognize the target relies on understanding the main structural features, including local structural dynamics. Furthermore, developing selective therapies and efficient vaccines against adaptive evolutionary patterns of the virus poses a significant challenge. This challenge is amplified due to the persistency of the pandemic and the estimation that SARS-CoV-2 might continue to circulate in the population with renewed outbreaks (Kissler, et al., 2020; Tse, et al., 2020; Ye, et al., 2020). Our analysis has laid the major inducible structural features of the SARS-CoV-2-RBD and propose a new potential therapeutic strategy to block viral entry. Overall, this study may be helpful in guiding the development and optimization of structure-based intervention strategies that target SARS-CoV-2.

## Acknowledgements

The authors would like to thank Haya and Nehemia Lemelbaum for their continuous generous support.

