## Supplementary Figure 1 for "Structural basis of SARS-CoV-2 spike protein induced by ACE2"

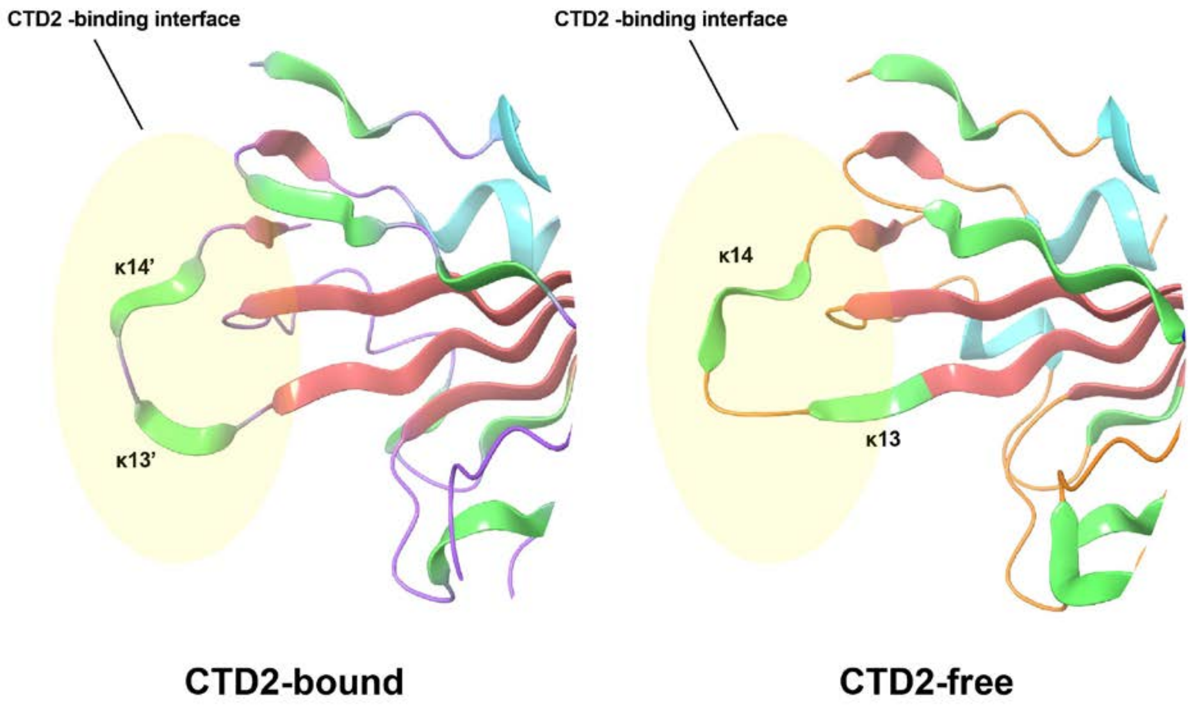


**Supplementary Figure 1.** Dissociation between the RBD and CTD2. The κ13/κ14 loop between residues 515 and 523 exhibits a conformational change between the unbound-closed (6VXX:B) and the bound (6M0J:E) states are shown as cartoon representation. Key contacts are labeled and shown as sticks. The secondary structural elements are shown as a cartoon. α-helices (cyan), β-strands (red), and κ-helices (green) are illustrated as ribbons.
